# The Orphan Kinesin PAKRP2 Achieves Processive Motility Via Noncanonical Stepping

**DOI:** 10.1101/430736

**Authors:** Allison M. Gicking, Pan Wang, Chun Liu, Keith J. Mickolajczyk, Lijun Guo, William O. Hancock, Weihong Qiu

## Abstract

PAKRP2 is an orphan kinesin in *Arabidopsis thaliana* that is thought to transport vesicles along phragmoplast microtubules for cell plate formation. Here, using single-molecule fluorescence microscopy, we show that PAKRP2 exhibits processive plus-end-directed motility on single microtubules as individual homodimers despite having an exceptionally long (32 residues) neck linker. Furthermore, using high-resolution nanoparticle tracking to visualize motor stepping dynamics, we find that PAKRP2 achieves processivity via a noncanonical stepping mechanism that includes small step sizes and frequent lateral steps to adjacent protofilaments. We propose that the small steps sizes are due to a transient intermediate step that involves a prolonged diffusional search of the tethered head due to its long neck linker. Despite this different stepping behavior, ATP is tightly coupled to each 8-nm step. Collectively, this study reveals PAKRP2 as the first orphan kinesin to demonstrate processive motility and broadens our understanding of the diverse kinesin stepping mechanisms.

## INTRODUCTION

Kinesins constitute a diverse superfamily of ATP-dependent, microtubule-based motor proteins that are known to participate in a variety of intracellular processes, such as microtubule organization and dynamics (1–5) and transport of cellular cargos (6, 7). Previous studies have revealed that some kinesins can move on microtubules by taking many consecutive steps before falling off, allowing them to transport organelles and protein complexes long distances (7–9). A specific example of cellular cargo transport occurs exclusively in plant cell division, which requires the construction of a cell plate at the division site. Plant kinesins hauling cell plate material walk on a microtubule-based structure called the phragmoplast, whose plus-ends are located at the developing cell wall (10, 11). In the model plant organism *Arabidopsis thaliana*, the phragmoplast-associated kinesin PAKRP2 is believed to transport Golgi vesicles to the phragmoplast midzone (12).

On the basis of phylogenetic analysis of the motor domains, the kinesin superfamily is divided into 14 subfamilies (kinesin-1 through kinesin-14) and an “orphan” (or ungrouped) family (13). PAKRP2 falls into the orphan family due to divergent structural features, such as a mutation in the conserved nucleotide binding site (12, 14). To date, no processive orphan kinesin has been reported, and many of the characterized orphan kinesins have mutations in the conserved residues essential for motility (15–17). Based on our current knowledge, PAKRP2’s predicted function of long-distance vesicle transport is contradictory to its classification as an orphan kinesin, but no investigation of its motility has ever been done. While processive motility isn’t necessarily required for organelle transport (18), it is a conserved feature of the prototypical transport kinesins, such as kinesin-1 and kinesin-2 (19, 20). Currently, it is not known if PAKRP2 is intrinsically processive or achieves transport of cargo via clustering of several diffusive motors (18, 21) or a non-motor microtubule binding domain that enhances the microtubule affinity of the motor domain via tethering (22, 23).

In this study, using a combination of single-molecule fluorescence microscopy, dark-field nanoparticle tracking, and solution ATPase assays, we show that PAKRP2 is an inherently processive kinesin and surprisingly, does not exhibit typical hand-over-hand stepping behavior on a single protofilament. Close examination of the PAKRP2 sequence revealed that it contains a long neck linker that contributes to its processive behavior and does not adversely affect the motor coupling. Overall, this study provides the first glimpse at the motility of an orphan kinesin and broadens current understanding of how kinesin structure and processivity are correlated.

## MATERIALS AND METHODS

### Molecular cloning, protein expression and purification

The full-length cDNA of PAKRP2 was codon-optimized for protein expression in *E. coli* and synthesized as two gBlocks (IDT). The construct, containing a C-terminal His-Tag, was integrated in a modified pET17b vector via isothermal assembly and verified by DNA sequencing. All of the truncation constructs were designed for this study except for K560AviC, which was created and characterized previously (24). For protein expression, plasmids were transformed into the BL21 Rosetta (DE3) competent cells (Novagen). Cells were grown at 37 ^o^C in TPM (containing 20g tryptone, 15g yeast extract, 8g NaCl, 2g Na_2_HPO_4_ and 1g KH_2_PO_4_ per 1 liter) supplemented with 50 *μ*g/ml ampicillin and 30 *μ*g/ml chloramphenicol. Expression was induced by cold shock on ice at OD_600_ = 0.8-1 with 0.1 mM IPTG, and incubation was continued for additional 14–17 hours at 20 ^o^C. Cell pellets were harvested by centrifugation at 5,500 x g for 10 minutes using a S-5.1 rotor (Beckman Coulter) and stored at −80 ^o^C prior to cell lysis.

To purify the His-tagged PAKRP2 and kinesin-1 chimera motors, cell pellets were re-suspended in the lysis buffer (50 mM sodium phosphate buffer, pH 7.2, 250 mM NaCl, 1 mM MgCl_2_, 0.5 mM ATP, 10 mM ß-mercaptoethanol, 20 mM imidazole, and 1 *μ*g/ml Leupeptin, 1 *μ*g/ml Pepstatin, 1 mM PMSF and 5 % glycerol), and lysed via sonication (Branson Sonifier 450). The cell lysate was then centrifuged at 21,000 × g for 35 minutes using a Ti-75 rotor (Beckman Coulter). The supernatant was incubated with Talon beads (Clontech) by end-to-end mixing at 4 ^o^C for 1 hour. The protein/beads slurry was then applied to a Poly-Prep column (Bio-Rad) and washed twice with 10 column volumes of wash buffer (50 mM sodium phosphate buffer, pH 7.2, 250 mM NaCl, 1 mM MgCl_2_, 0.1 mM ATP, 10 mM ß-mercaptoethanol, 20 mM imidazole, and 1 *μ*g/ml Leupeptin, 1 *μ*g/ml Pepstatin, 1 mM PMSF and 5% glycerol). The protein was eluted with 5 column volumes of elution buffer (50 mM sodium phosphate buffer, pH 7.2, 250 mM NaCl, 1 mM MgCl_2_, 0.5 mM ATP, 10 mM ß-mercaptoethanol, 250 mM imidazole and 5 % glycerol). The eluted protein was buffer-exchanged with a PD-10 column into storage buffer (BRB80, 0.5 mM ATP, 100 mM KCl and 5% glycerol), flash frozen in liquid nitrogen, and stored at −80 ^o^C.

### Polarity-marked microtubules

To make polarity-marked GMPCPP microtubules, a dim bovine tubulin mix (containing 17 *μ*M unlabeled tubulin, 17 *μ*M biotinylated tubulin, and 0.8 *μ*M HiLyte 647-tubulin) was first incubated in BRB80 with 0.5 mM GMPCPP (Jena Bioscience) at 37° C overnight to make dim microtubules, and then centrifuged at 250,000 × g for 7 minutes at 37° C in a TLA100 rotor (Beckman Coulter). The pellet was re-suspended in a bright bovine tubulin mix (containing 7.5 *μ*M unlabeled tubulin, 4 *μ*M HiLyte 647-tubulin, and 15 *μ*M NEM-tubulin) in BRB80 with 2 mM GMPCPP and incubated at 37° C for additional 15 minutes to cap the plus-end of the dim microtubules. The resulting polarity-marked track microtubules were pelleted at 20,000 × g for 7 minutes at 37° C in the TLA100 rotor (Beckman Coulter), and finally re-suspended in BRB80 with 40 *μ*M taxol.

### Total internal reflection fluorescence (TIRF) microscopy

All time-lapse imaging assays were performed at room temperature (22–23 °C) using the Axio Observer Z1 objective-type TIRF microscope (Zeiss) equipped with a 100x 1.46 NA oil-immersion objective and a back-thinned electron multiplier CCD camera (Photometrics). All microscope coverslips were functionalized with biotin-PEG as previously described (25) to reduce nonspecific surface absorption of molecules. All time-lapse imaging experiments in this study used flow chambers that were made by attaching a coverslip to a microscope glass slide by double-sided tape.

### Single-molecule motility assay

For single-molecule motility experiments, the chamber was perfused with 0.5 mg/ml Streptavidin to immobilize the taxol-stabilized polarity-marked HiLyte-647/Biotin-labeled microtubules. After removing unbound microtubules by washing the chamber with five-chamber volumes of BRB12 supplemented with 20 *μΜ* taxol, the chamber was perfused with a BRB80-based (or BRB50 for Kin1_NLswap) motility mixture containing diluted motors, 1 mM ATP, 25 *μΜ* taxol, 1.3 mg/ml casein and an oxygen scavenger system based on glucose oxidase/catalase (26). Time-lapse image sequences were recorded at 1 frame per 2 seconds with an exposure time of 200 ms for a duration of up to 10 minutes. Kymographs were generated and analyzed in ImageJ (NIH) for obtaining the velocity and run length information of individual PAKRP2 motors. Reported velocities are the peak of a Gaussian fit to the data, and associated errors are the standard deviations (SD). Characteristic run lengths were calculated by fitting an exponential cumulative distribution function to the data, and creating a bootstrap distribution to find the mean (n = 5000). The mean was then corrected for filament length using the procedure outlined in (27), where the characteristic filament length is the average length of all microtubules used in the analysis. Reported errors are the 95% confidence intervals (CI) of the bootstrap distributions.

### Single-molecule photobleaching assay

For single-molecule photobleaching assays, PAKRP2 molecules were immobilized, in the absence of ATP, on taxol-stabilized polarity-marked HiLyte-647/Biotin-labeled microtubules in BRB80 with 20 *μ*M taxol and 1.3 mg/ml casein. Time-lapse image sequences were continuously recorded with an exposure time of 100 ms until the field of view was completely bleached of fluorescence signal. The number of photobleaching steps of individual PAKRP2 motors was obtained by tracking the fluorescence intensity in ImageJ (NIH).

### Total internal reflection dark-field microscopy assays

For single-molecule tracking experiments, we used a custom-built total internal reflection dark-field microscope as previously described (28). All experiments were carried out at 22–23 °C. PAKRP2 motors were prepared with a biotinylated C-terminal Avitag or biotinylated GFP binding protein (29), and mixed with 30-nm diameter, Streptavidin-coated gold nanoparticles. The motor was added to the gold particles at the lowest molar ratio that produced landing events (4:1 for PAKRP2 and 3:1 for K560AviC). Taxol-stabilized GDP microtubules were attached to the glass coverslip via a kinesin rigor mutant as previously described (24). High-resolution position vs. time traces were obtained from 100 frame/s (or 1000 frame/s for kinesin-1) movies by fitting the point spread function with a 2D Gaussian using Fiesta software (30). The x and y trajectories were rotated to minimize the standard deviation in the x-direction, resulting in a y-axis that is aligned with the microtubule axis.

### Step size determination

The on-axis step sizes were determined by a t-test algorithm (31), for the y-displacement vs time traces. Only the traces with a standard deviation of < 3 nm from the step plateau were chosen for analysis. All reported step sizes are the mean ± standard deviation of a single Gaussian fit to the forward steps.

### ATPase Assay

ATPase assays were carried out by an enzyme-coupled assay protocol adapted from a previous study (32). Assays used 25 nM active dimeric PAKRP2(560), and 5 nM active dimeric Kin1_NLswap, where activity was determined by assessing exchange of mantADP, as previously described (33). Hydrolysis rates at each microtubule concentration (GDP taxol-stabilized) were estimated by a linear fit to steady-state absorbance decreases at 340 nm, as previously described (33). The k_cat_ and K_M_ were determined by performing a least-squares fit of the ATPase vs. tubulin concentration curve to the Michaelis-Menten equation.

## RESULTS

### PAKRP2 is a Processive Kinesin with a Long Neck Linker

PAKRP2 consists of an N-terminal motor domain followed by a neck linker, a coiled-coil central stalk, and an uncharacterized C-terminal tail. Based on previous studies that show that the neck linker is an important component of kinesin processivity (34), and the fact that PAKRP2 is an orphan kinesin (14, 35), we first wanted to determine if there were any structural divergences in the neck linker region. Two coiled-coil prediction programs, COILs (36) and MARCOIL (37), both placed the likely start of the alpha-7 helix between residues 394 and 407 (Fig. 1 *A* and *B*). Examination of the sequence within that range places the first logical heptad repeat of the coiled-coil domain at residue 397, a hydrophobic methionine, followed by negatively charged aspartic and glutamic acids (Fig. 1 *C*). Thus, we conclude that the neck linker of PAKRP2 contains 32 residues, considerably longer than the neck linkers of other kinesins, which typically contain 14–18 residues (38).

**Figure 1:**
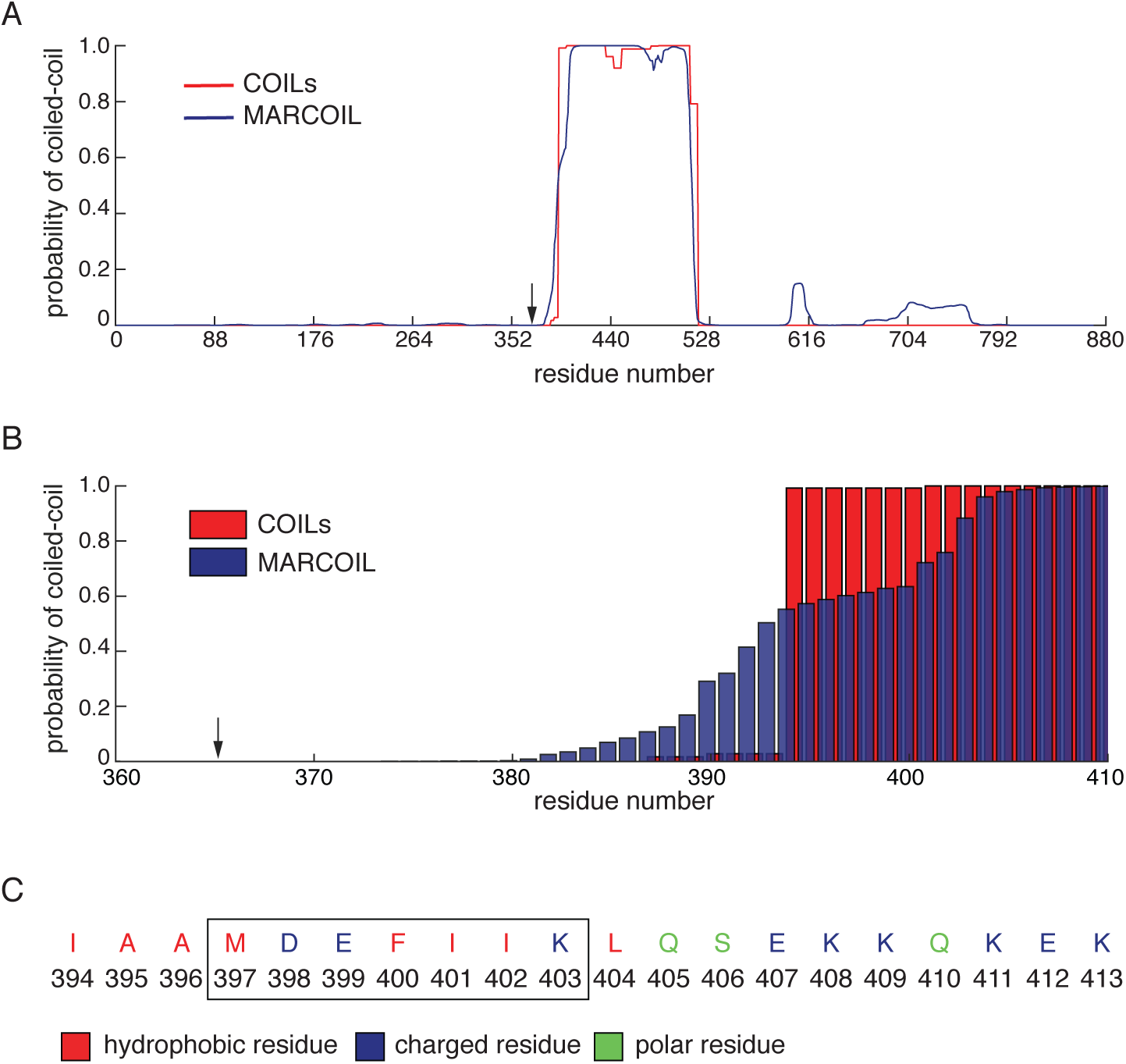
PAKRP2 contains a long neck linker. (A) The coiled-coil profiles of full-length PAKRP2 predicted by COILs (red) and MARCOIL (blue). Black arrow denotes the end of the motor domain. (B) Zoomed-in coiled-coil profiles of residues just before the coiled-coil region. Black arrow denotes the end of the motor domain. (C) Sequence of residues between 394 and 414 to show first heptad repeat starting at residue 397 and ending at residue 403.

Several studies on kinesin-1 and kinesin-2 demonstrate that increasing the neck linker length can lead to disruptions in the stepping mechanism, such as a decrease in run length or an increase in futile ATP hydrolysis cycles (39–42). To determine whether PAKRP2 is a processive microtubule motor, we engineered PAKRP2(FL), a recombinant full-length PAKRP2 with a C-terminal GFP (Fig. 2 *A* and *B*). We used a single-molecule motility assay to visualize the movement of PAKRP2(FL) on polarity-marked microtubules (Fig. 2 *C* and *D*; Supplementary Movies 1 and 2). The assay showed that individual PAKRP2(FL) molecules moved processively toward microtubule plus-ends with a mean velocity of 65 ± 16 nm s^−1^ (mean ± SD, n = 271; Fig. 2 *E*) and a run length of 3.56 ± 0.27 *μ*m (mean ± 95% CI, n = 271, Fig. 2 *F*). The reported run lengths and the associated error are the mean and 95% confidence intervals of the bootstrap distribution. It is well established that nonprocessive kinesin motors can achieve processive motility by clustering to form multi-motor ensembles (18, 21). We thus performed single-molecule photobleaching to determine the oligomerization of PAKRP2(FL). Similar to other dimeric kinesins (18), PAKRP2(FL) was predominantly photobleached in one or two steps (Fig. S1, *A* and *B*). These results show that PAKRP2(FL) contains the ability to exhibit processive plus-end-directed motility on single microtubules as a homodimer.

**Figure 2:**
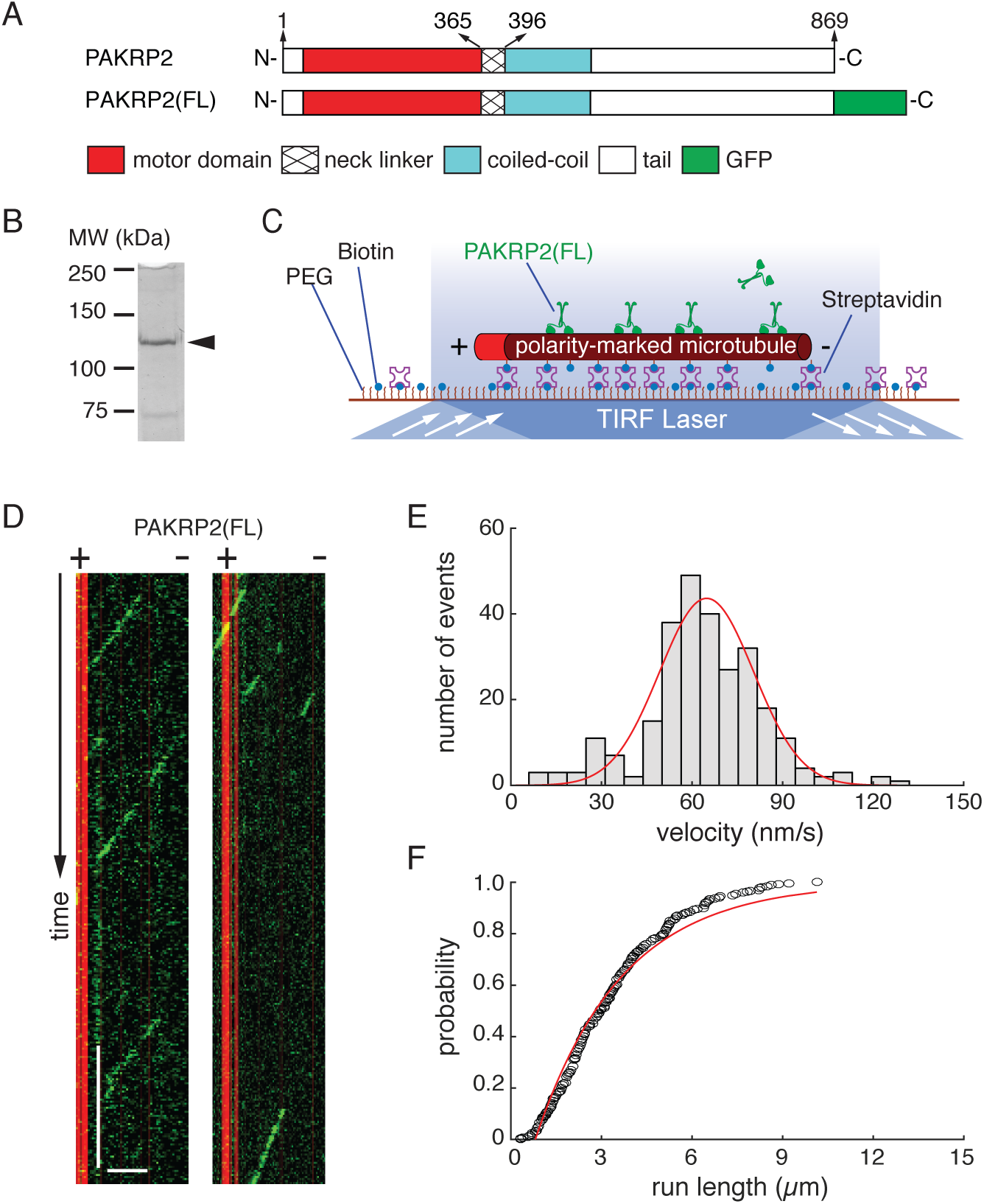
Full-length PAKRP2 is a processive plus-end-directed kinesin. (A) Schematic diagrams of full-length PAKRP2 and PAKRP2(FL). (B) SDS-PAGE gel of PAKRP2(FL). Arrowhead indicates the expected band of PAKRP2(FL), MW = 126 kDa. (C) Schematic diagram of the single-molecule motility assay. (D) Representative kymographs of individual PAKRP2(FL) molecules moving processively towards the plus-ends of single microtubules. Scale bars: 2 minutes (vertical); 5 *μ*m (horizontal). (E) Velocity histogram of PAKRP2(FL). Red line corresponds to a Gaussian fit with a mean value of 65 ± 16 nm s^−1^ (n = 271). (F) Cumulative distribution of PAKRP2(FL) run length with a characteristic run length of 3.56 ± 0.27 *μ*m (n = 271). Black circles correspond to experimental data and the red line corresponds to an exponential cdf fit.

Some kinesins are known to achieve processive motility or to gain enhanced processivity via non-motor microtubule-binding domains (22, 23, 43). To test whether PAKRP2 processivity depends on any domain beyond the head and neck linker domains, we made two additional constructs: PAKRP2(560), a truncation of the full-length at residue 560, and PAKRP2(LZ), a minimal dimer of PAKRP2 containing the motor domain and the neck linker and dimerized through a leucine zipper (Fig. 3 *A-C*). Single-molecule photobleaching experiments confirmed that PAKRP2(560) and PAKRP2(LZ) both exist predominantly as individual homodimers (Fig. S1 *C*-*F*), and exhibited processive plus-end-directed motility on single microtubules (Fig. 3 *D*, Supplementary Movies 3 and 4). The velocities of PAKRP2(560) and PAKRP2(LZ) were within error of PAKRP2(FL) (Fig. 3 *E*, Fig. S2, *A* and *C*). Run lengths of PAKRP2(560) and PAKRP2(LZ) were determined to be 3.35 ± 0.29 *μ*m (n = 266, Fig. 3 *E*, Fig. S2 *B*) and 2.67 ± 0.25 *μ*m (n = 333, Fig. 3 *E*, Fig. S2 *D*), respectively. While the PAKRP2(560) run length was within 5% of the wild type run length, the more significant decrease in run length of the PAKRP2(LZ) construct could indicate the native coiled-coils play a role in motility. Overall, these data show that PAKRP2 processivity is largely encoded in the region containing the motor domain and the neck linker.

**Figure 3:**
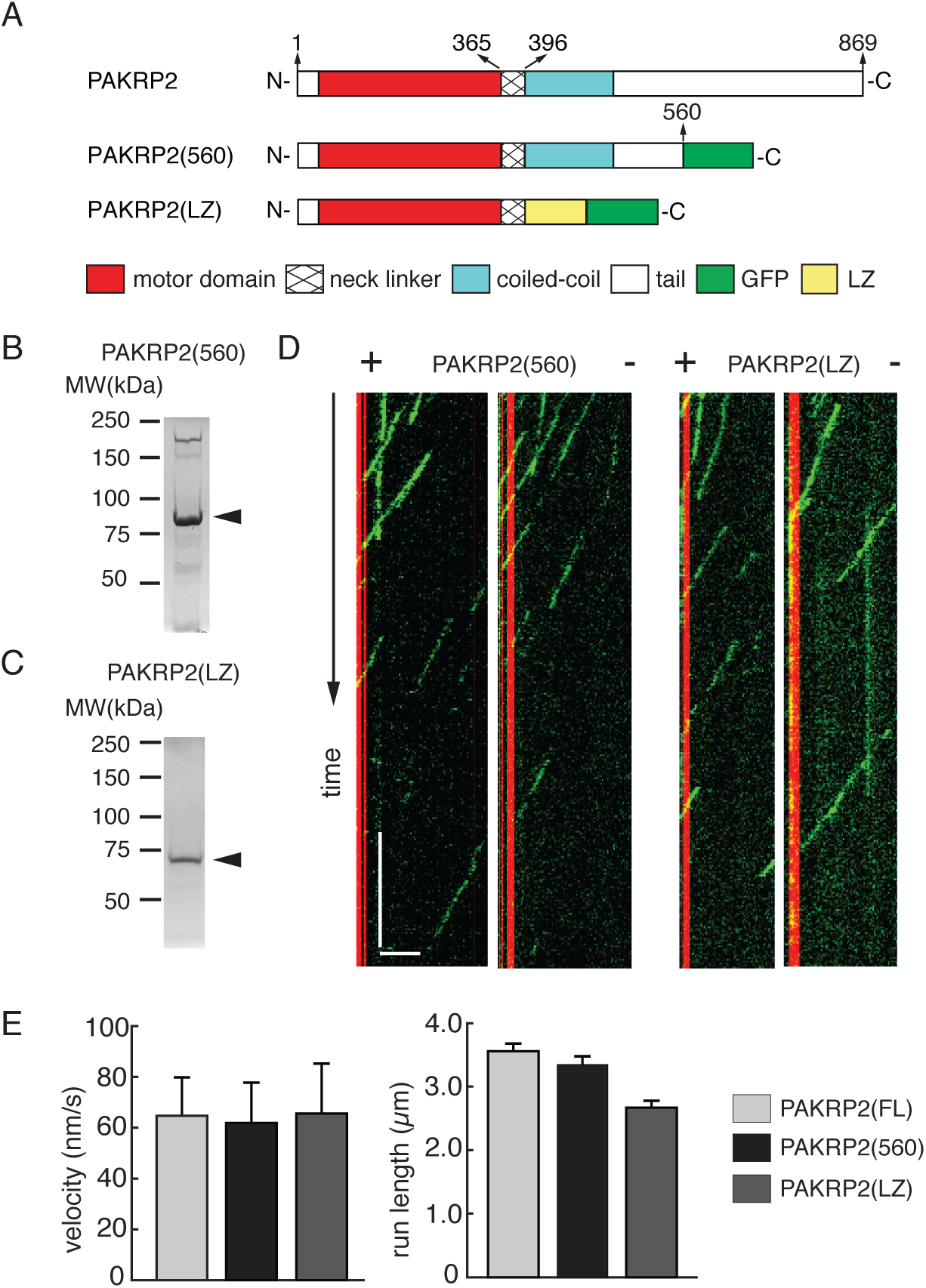
PAKRP2 processivity does not require C-terminal tail or stalk domain. (A) Schematic diagram of full-length PAKRP2, PAKRP2(560) and PAKRP2(LZ). (B) SDS-PAGE gel for PAKRP2(560). Arrowhead indicates the expected band of PAKRP2(FL). MW = 91 kDa. (C) SDS-PAGE gel for PAKRP2(LZ). Arrowhead indicates the expected band of PAKRP2(FL). MW = 76 kDa. (D) Representative kymographs of individual PAKRP2(560) and PAKRP2(LZ) molecules on single microtubules. Scale bars: 2 minutes (vertical); 5 *μ*m (horizontal). (E) Bar graphs showing the relative velocities and run lengths of PAKRP2(FL), PAKRP2(560) and PAKRP2(LZ) with corresponding errors.

### PAKRP2 Frequently Moves Laterally

Based on the knowledge that PAKRP2 is inherently processive despite having a long neck linker, we next investigated its stepping behavior to determine whether and how it may differ from that of kinesin-1. We attached a 30-nm gold particle to the C-terminus of PAKRP2(560) and observed the center-of-mass motion via dark-field nanoparticle tracking (Fig. 4 *A*) (44). Gold nanoparticle attachment did not significantly affect the motor activity, as the single-molecule velocity of nanoparticle-labeled motors on taxol-stabilized GDP microtubules was within 20% of the motors without nanoparticles in the same experimental conditions (Fig. S3). It should also be noted that the velocity of PAKRP2(560) without a conjugated gold nanoparticle on GDP microtubules was 36% slower than its velocity without a conjugated gold nanoparticle on GMPCPP microtubules (Fig. S3). This suggests that PAKRP2(560) is sensitive to nucleotide-dependent structural changes of the microtubule lattice (45), which has also been observed in the motility of kinesin-1 (46, 47).

**Figure 4:**
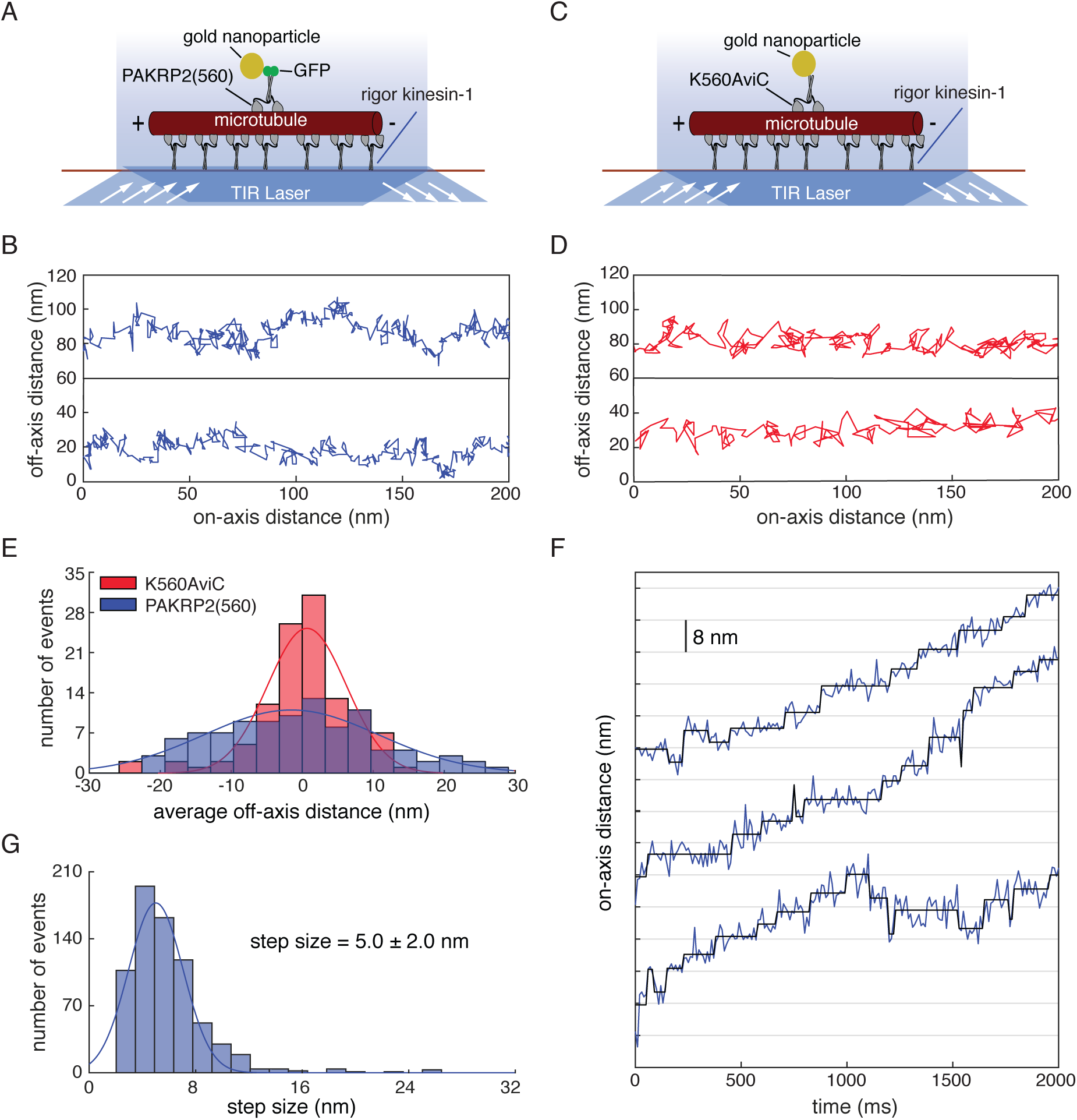
PAKRP2 takes frequent lateral steps. (A) Schematic diagram of the gold nanoparticle placement on the C-terminus of PAKRP2(560) (B) Sample x-y traces for PAKRP2(560), where the x-axis corresponds to the microtubule axis and the y-axis corresponds to lateral movement. (C) Schematic diagram of the gold nanoparticle placement on the C-terminus of K560AviC. (D) Sample x-y traces for PAKRP2(560), where the x-axis corresponds to the microtubule axis and the y-axis corresponds to lateral movement. (E) Off-axis distance distributions for PAKRP2(560) and K560AviC. Red and blue lines correspond to a Gaussian fit to the data. (F) A representative on-axis distance vs. time plot. Raw data is shown as blue lines and steps detected by the t-test step finding algorithm are shown in black. Data was acquired at 1 mM ATP every 10 ms. (G) Step size histogram of PAKRP2(560) fit to a single Gaussian (n = 958).

When imaged at high resolution, PAKRP2(560) appeared to take frequent and sequential lateral steps both to the left and right (Fig. 4 *B*). To rule out that this behavior was due to surface binding of the gold nanoparticle or some other artifact of the assay, we used kinesin-1 with a gold nanoparticle on the C-terminus as a control (K560AviC, Fig. 4 *C*), since kinesin-1 has been demonstrated to walk on single protofilament unless it is navigating a roadblock (48, 49). The different lateral stepping characteristics of PAKRP2(560) with a gold nanoparticle on the C-terminus can be seen clearly when compared to K560AviC (Fig. 4 *D*). To quantify the lateral stepping behavior, a distribution of the lateral displacement per 40 nm of on-axis displacement was generated for both motors (Fig. 4 *E*). The standard deviations for the Gaussian fits to the histograms of average off-axis displacements were 12.3 nm (n = 99) and 5.1 nm (n = 113) for PAKRP2 and kinesin-1, respectively, confirming that PAKRP2 steps laterally more than kinesin-1. Interestingly, the respective means were −1.5 nm and 0.7 nm, indicating that neither motor has a significant off-axis directional bias.

Canonical processive kinesins are known to take sequential 8-nm center-of-mass steps for the duration of their run lengths, due to the periodicity of tubulin binding sites on a single protofilament (50–52). Using point-spread-function fitting to the gold nanoparticle position and a t-test step-finding algorithm (Fig. 4 *F*), we measured an on-axis, center-of-mass step size of 5.0 ± 2.0 nm for PAKRP2(560) (mean ± SD, n = 958, Fig. 4 *G*). In comparison, the kinesin-1 control displayed an on-axis, center-of-mass step size of 8.0 ± 3.0 nm (mean ± SD, Fig. S4 *A* and *B*), consistent with previous results (49, 51). Thus, PAKRP2 stepping is distinct from that of kinesin-1, in both the lateral movement and the average step size.

### PAKRP2 Takes Intermediate Steps

To better understand the individual head dynamics during PAKRP2 stepping and to confirm the small step size seen in the center-of-mass data, we attached the gold nanoparticle to one motor domain (head) of PAKRP2(560) via an N-terminal Avi-tag (Fig. 5 *A*). Similar to the center-of-mass construct, the head-tagged motor stepped processively along the microtubule with clearly observable steps (Fig. 5 *B*). Using the t-test step-finding algorithm, we measured an average step size of 7.6 ± 3.4 nm (mean ± SD, n = 230, Fig. 5 *C*). This value is larger than the center-of-mass step size, as expected, but is half of the 16.4 nm expected for canonical hand-over-hand stepping (39, 53). There are two plausible explanations for a step size smaller than the distance between successive binding sites on a single protofilament: the labeled head is binding to an adjacent protofilament or the steps represent transient intermediates in which the labeled head is between binding sites. For instance, Stepp *et al*. demonstrated that head-labeled kinesin-2 motors take 13-nm steps on axonemes, compared to 16.4-nm steps on single microtubules (54). This discrepancy was explained by 50% of the steps landing on an adjacent site 8.2 nm away. In contrast, high-resolution nanoparticle tracking revealed that the motor domain of kinesin-1 takes intermediate steps, or substeps, at saturating ATP conditions that result in an average step size of 8.2 nm (24).

**Figure 5:**
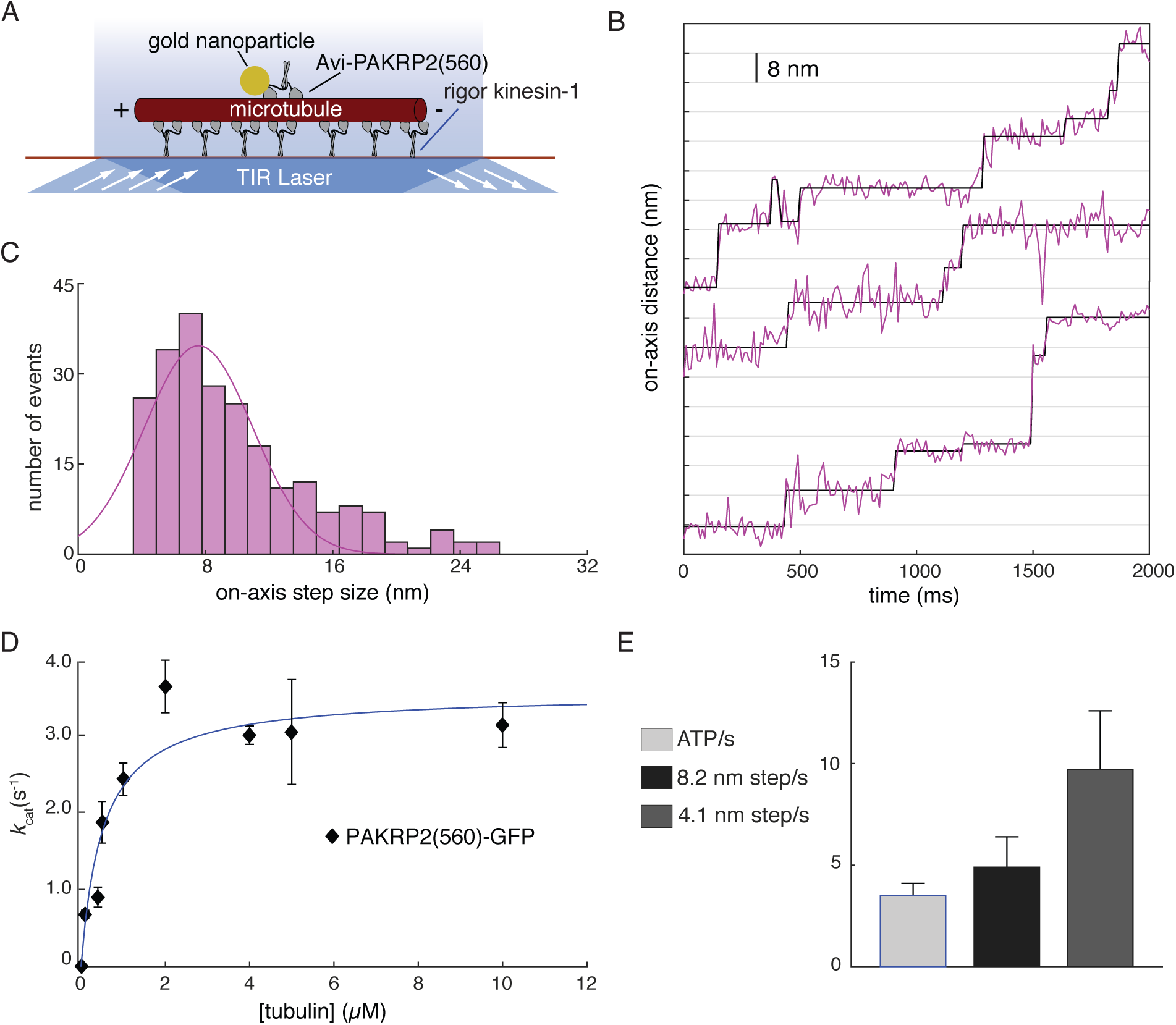
PAKRP2 takes intermediate steps. (A) Schematic diagram of the gold nanoparticle placement on the N-terminal motor domain of Avi-PAKRP2(560). (B) A representative on-axis distance vs. time plot. Raw data are shown as purple lines and steps detected by the t-test step finding algorithm are shown in black. Data were acquired at 1 mM ATP every 10 ms. (C) Step size histogram of Avi-PAKRP2(560) fit to a single Gaussian (n = 230). (D) The ATPase rate per dimer for PAKRP2(560) as a function of tubulin concentration. Data correspond to the mean ± SE (n = 3 or 5 determinations per point). The solid line represents a fit to the Michaelis-Menten equation, giving a maximum k_cat_ of 3.5 ± 0.6 s^−1^. (E) Comparison of the ATP hydrolysis rate per dimer and the stepping rate per dimer assuming different step sizes. Stepping rates were calculated from the division of the center-of-mass velocity by the center-of-mass step size.

In order to resolve whether the PAKRP2 motor domains are taking 8.2-nm steps to adjacent protofilaments or pausing midway through 16.4-nm steps, we measured the ATPase activity of PAKRP2. Canonical steppers are known to take one step per ATP molecule hydrolyzed, as ATP binding initiates the power stroke (55, 56). It is important to remember here that the displacement of the motor’s center-of-mass is approximately half the displacement of one motor head if the second head remains bound to the microtubule. Based on the measured velocity of 40 ± 12 nm s^−1^ on GDP taxol-stabilized microtubules (Fig S3), 8.2-nm steps of the center-of-mass correspond to a stepping rate of 4.9 s^−1^ and 4.1-nm steps correspond to a stepping rate of 9.8 s^−1^. In the ATPase assay the k_cat_ for a PAKRP2(560) dimer was 3.5 ± 0.6 ATP/s (Fig. 5 *D*), which is close to the rate for 8.2 nm center-of-mass steps (Fig. 5 *E*) and not consistent with the motor hydrolyzing multiple ATP per step. Therefore, we conclude that PAKRP2 stepping includes an intermediate step, with the final position of the head being ~16.4 nm from the starting position. This result also necessitates that PAKRP2 uses one ATP per step, despite having a long neck linker, which is contrary to previous studies on kinesin-1 in which long neck linkers lead to uncoupling of the ATPase from stepping (39, 42).

### PAKRP2 Neck Linker Disrupts Kinesin-1 Stepping

The finding that PAKRP2 contains a long neck linker domain yet retains tight coupling between its ATPase and stepping activities raises the possibility that sequence-specific structural features in its long neck linker contribute to the coupling. To test whether the PAKRP2 neck linker confers tight coupling to other motors, we designed a kinesin-1 chimera, Kin1_NLswap, in which the native 14 residue neck linker was replaced by the 32 residue PAKRP2 neck linker (Fig. 6 *A*, S5 *A*). A photobleaching assay confirmed that this construct is a homodimer in solution (Fig. S5, *B* and *C*). From kymograph analysis (Supplementary Movie 5), the single molecule velocity of Kin1_NLswap was 103 ± 20 nm s^−1^ (n = 172, Fig. 6 *B*), and the run length was 1.64 ± 0.19 *μ*m (n = 172, Fig. 6 *C*). Under identical conditions, the velocity and run length of wild-type kinesin-1 are 670 ± 70 nm s^−1^ and 1.21 ± 0.16 *μ*m, respectively (44). Thus, swapping the PAKRP2 neck linker into kinesin-1 significantly disrupts the stepping rate but does not substantially alter motor processivity.

**Figure 6:**
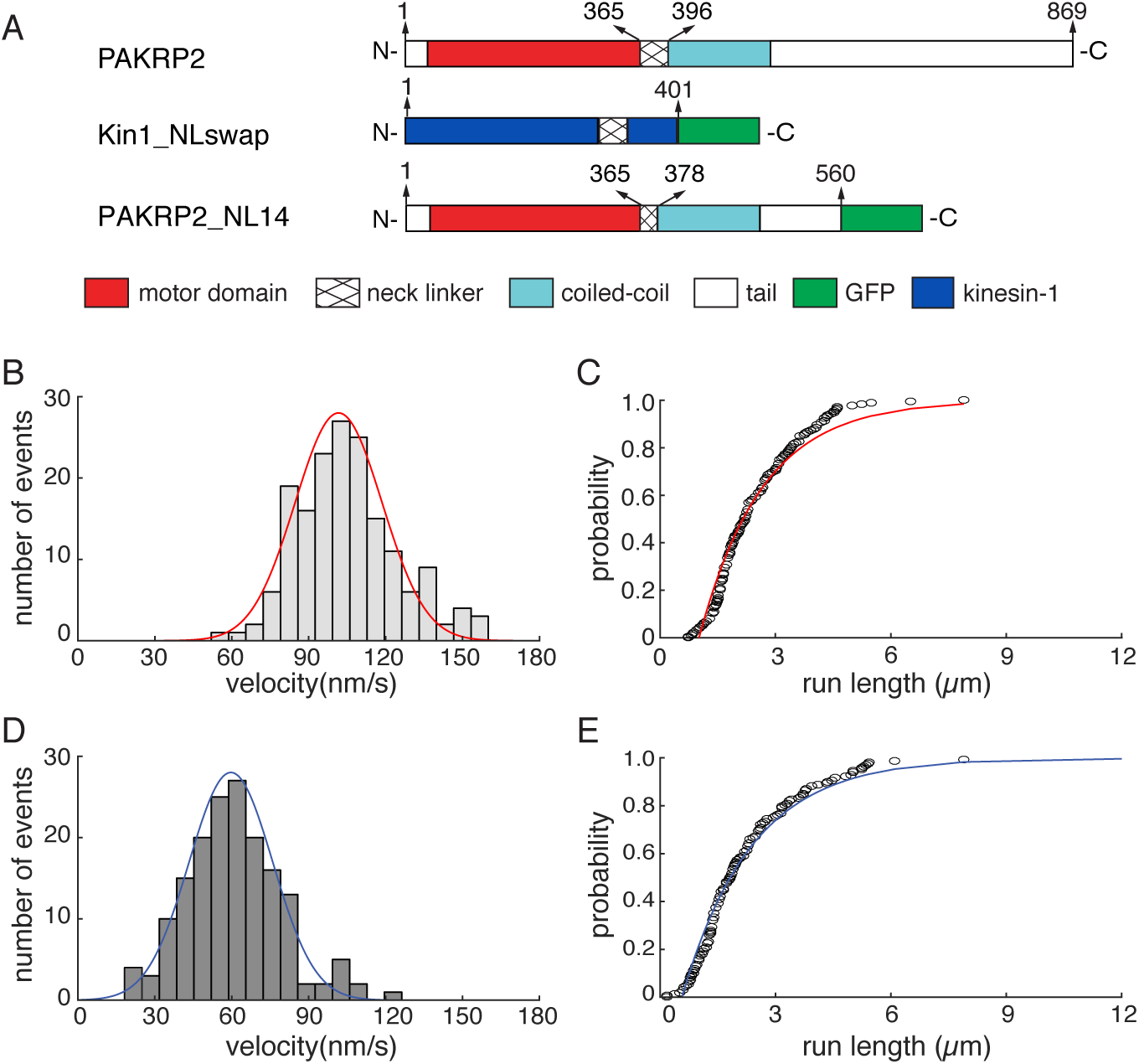
PAKRP2 neck linker affects processivity. (A) Schematic diagram of Kin1_NLswap and PAKRP2_NL14. (B) Velocity histogram of Kin1_NLswap. Red line corresponds to a Gaussian fit with a mean of 103 ± 20 nm s^−1^ (n = 172). (C) Run length histogram of Kin1_NLswap. Red line corresponds to an exponential fit with a characteristic run length of 1.64 ± 0.19 *μ*m (n = 172). (D) Velocity histogram of PAKRP2_NL14. Blue line corresponds to a Gaussian fit with a mean of 59 ± 19 nm s^−1^ (n = 168). (E) Run length histogram of PAKRP2_NL14. Blue line corresponds to an exponential fit with a characteristic run length of 1.96 ± 0.27 *μ*m (n = 168).

In principle, this slower velocity could result from either slowing of the ATPase cycle or uncoupling of the hydrolysis cycle from the stepping cycle. To test between these possible mechanisms, we measured the microtubule-stimulated ATPase of Kin1_NLswap. The k_cat_ of the Kin1_NLswap was 121 ± 7 s^−1^ (Fig. S6), which is significantly higher than the center-of-mass stepping rate of ~12 s^−1^, assuming it takes 8.2-nm steps. However, even if the steps were 4.1-nm, the stepping rate of ~25 s^−1^ would still be considerably lower than the ATP hydrolysis rate. Thus, replacing the kinesin-1 neck linker with the longer PAKRP2 neck linker led to uncoupling of the kinesin-1 from its stepping activity. It follows that, although PAKRP2 maintains tight mechanochemical coupling despite having a long neck linker, tight coupling results from features of the motor catalytic domain rather than specific structural features of the neck linker domain.

### PAKRP2 Neck Linker Enhances Its Processivity

Given that the PAKRP2 neck linker greatly disrupts the stepping cycle of kinesin-1, it could also negatively impact the stepping cycle of PAKRP2. To determine if the neck linker extension affects the processivity of PAKRP2, we made a mutant, PAKRP2_NL14, in which neck linker residues beyond the first fourteen amino acids in the sequence were deleted (Fig. 6 *A* and S5 *D*). The construct was also confirmed to be a homodimer (Fig. S5, *E* and *F*). In contrast to the change in velocity resulting from swapping the neck linker into Kinesin-1, shortening the PAKRP2 neck linker did not affect the stepping rate. The velocity of PAKRP2_NL14 on GMPCPP microtubules was determined to be 59 ± 19 nm s^−1^ (n = 168, Fig. 6 *D*), within 10% of wild-type PAKRP2 (Fig. 2 *E*). Additionally, single-molecule motility assays showed that the neck-shortened motor maintains processivity, but the run length is decreased to 1.96 ± 0.27 *μ*m (n = 168, Fig. 6 *E*), which is 45% less than wild-type (Supplementary Movie 6). The deleted region of the neck linker has a neutral charge, making it unlikely that the reduction in run length is due to weaker electrostatic interactions between the motor domain and the microtubule (57). Therefore, unlike in other kinesins where longer neck linkers reduce inter-head coordination leading to decreased processivity, the long neck linker of PAKRP2 contributes to its substantial processivity.

## DISCUSSION

In this study, we have demonstrated that the orphan kinesin PAKRP2 is inherently processive, which sets it apart from all other characterized orphan kinesins but is consistent with its putative role in vesicle transport (12). Like other orphan kinesins (17, 58–60), PAKRP2 has a divergent nucleotide-binding motif and hydrolyzes ATP more than 10-fold slower than dimeric kinesin-1 (61, 62). It also contains a 32-residue neck linker that is much longer than most other members of the kinesin superfamily (38). Despite these variations in structural features that are integral to motility (40, 63), PAKRP2 is more processive than kinesin-1, and it achieves this processivity via a noncanonical stepping mechanism that includes a long one-head-bound intermediate state and frequent lateral steps.

In the kinesin-1 motility cycle, ATP turnover is tightly coupled to its stepping activity (55, 56, 64). This coupling is thought to be facilitated by conformational changes in the neck linker, which transmits tension between the two heads (41, 42). Evidence for this tension-based mechanism can be seen in the uncoupling of the heads in kinesin-1 with neck linker insertions, where in some cases the ATP turnover rate becomes much higher than the stepping rate (39, 42), and in some cases the stepping rate slows relative to wild-type (39, 40). Our study confirms this behavior in kinesin-1, as the ATP hydrolysis rate of Kin1_NLswap is much higher than the predicted stepping rate (Fig. S6). In contrast, PAKRP2 maintains tight mechanochemical coupling despite its long neck linker (Fig. 5 *E*), suggesting that the mechanism for motor coupling is different than in other kinesins. Consistent with this, shortening the PAKRP2 neck linker had no effect on the stepping rate but significantly decreased the run length (Fig. 6 *D* and *E*). These results suggest that the PAKRP2 motor domain and long neck linker have coevolved to achieve tight mechanochemical coupling through an alternative mechanism than the one that has been determined for kinesin-1.

High-resolution particle tracking of PAKRP2 revealed tail displacements of 5.0 nm (Fig. 4 *G*), which is considerably smaller than the 8.2-nm tubulin periodicity, and motor domain displacements of 7.6 nm (Fig. 5 *C*), which is considerably smaller than the 16.4 nm periodicity expected from a classical hand-over-hand mechanism (39, 42, 53). The small step size in the PAKRP2 head-labeled traces is characteristic of an inchworm-like stepping model whereby the two motor domains walk on adjacent protofilaments, such as the stepping observed in dynein motility (26, 65). Although dynein is capable of taking coordinated hand-over-hand steps, it has also been observed to be a stochastic stepper, wherein each head hydrolyzes ATP and moves independently of the other (26, 65). Inchworm-like stepping in this case differs from the originally proposed inchworm model in that it involves two catalytically active motor domains (66, 67), and therefore two ATP molecules are burned per 8.2 nm step. This type of stepping in dynein is facilitated by the flexibility between the two heads (68), making it a plausible mode of motion for a kinesin with a long neck linker. However, the ATPase data that show PAKRP2 burns one ATP molecule per 8.2 nm step oppose this explanation for the unusual step sizes seen in PAKRP2 (Fig. 5 *E*). This is not to say that PAKRP2 does not step on adjacent protofilaments, which was clearly shown in the center-of-mass data, but that the step does not end at a binding site 8.2 nm away.

An alternative explanation for the small PAKRP2 step sizes is that particle tracking is detecting an intermediate step (or substep) of the tethered motor head before it reaches the next binding site. A 2015 study on kinesin-1 labeled on one motor domain at saturating ATP observed 8.2-nm substeps that were attributed to a transient one-head-bound state following ATP binding and preceding ATP hydrolysis (24). In the present study, detection of these intermediate states by particle tracking could be facilitated by the slow stepping rate of PAKRP2, which is ~20-fold slower than kinesin-1. Step sizes smaller than 16.4 nm have been observed before in kinesin-1 mutants with long neck linkers, but they were not attributed to substeps (39). In support of a transient one-head-bound state, a recent study demonstrated that the duration of the one head-bound state can be increased by increasing the neck linker length in kinesin-1 and kinesin-2 motors, an effect that may result from an increase in the area of diffusional search taken by the tethered head before binding (44). Taken together, the PAKRP2 data align best with a substep model, though the exact mechanism of this substep may be different than what is seen in kinesin-1. In canonical processive steppers, such as kinesin-1 and kinesin-2, an increase in the one head-bound state duration leads to a higher probability that the bound head detaches before the trailing head binds, and consequently, causes a reduction in processivity of the motor (44). In contrast, shortening the neck linker of PAKRP2, which presumably decreases the one-head-bound state, resulted in reduced processivity (Fig. 6 *E*).

Another example of a kinesin with an unusually long neck linker is Zen4, a member of the kinesin-6 family that plays a role in microtubule organization during cytokinesis (69). The neck linker of Zen4 is 75 residues long and includes a binding site for GTPase activing proteins (70), but does not prevent the motor from processively stepping along microtubules (71). The crystal structure of the Zen4 motor domain in a nucleotide-free state revealed that the initial segment of the neck linker docked in a backward conformation; this conformation is thought to relieve inter-head tension and allow for more stability in the two head-bound state (71). Zen4 functions primarily in crosslinking microtubules, so the long neck linker, coupled with backward docking in the two head-bound state, could potentially allow both motor heads to remain bound for long periods of time. However, despite both motors having long neck linker domains, PAKRP2 does not contain the “arginine gate” that facilitates the backward docking of the neck linker, and it is not clear how stabilization of the two head-bound state would benefit a transport motor. Thus, parallels that can be drawn between Zen4 and PAKRP2 are limited.

Although long neck linkers have primarily been viewed as a disadvantage for processivity (20, 38–40), there is evidence to suggest that they are an advantage for obstacle avoidance. Kinesin-2, which contains a 17-residue neck linker, has been shown to step laterally to adjacent protofilaments (48) and to be less affected than kinesin-1 by the addition of roadblocks, such as tau protein or rigor kinesin-1 motors on the microtubule track (72, 73). These studies propose that a long neck linker makes kinesin-2 sufficiently flexible to step to many of the adjacent binding sites, and less likely to dissociate from the microtubule at a roadblock. In the cell, microtubules are decorated with microtubule-associated proteins (MAPs) that could potentially block the paths of processive motors (74–77), and so this obstacle avoidance could provide a selective advantage in maximizing transport. Given that PAKRP2 is thought to transport material on the phragmoplast microtubules, it seems likely that it encounters MAPs and side stepping would be a useful feature. As a final point, consecutive side-steps have also been observed for kinesin-8, which achieves superprocessivity despite having a long neck linker domain (23, 73, 78, 79); thus side-stepping does not necessarily correlate with decreased processivity.

Based on the data presented here, we propose that PAKRP2 steps via a hand-over-hand mechanism that includes a transient intermediate state in which one head is bound to the microtubule, and that the motor takes frequent lateral steps to adjacent protofilaments. We propose that this stepping behavior results from the long neck linker domain, however the tight mechanochemical coupling also suggests that the neck linker and catalytic core of the motor have coevolved to achieve processivity using a stepping mechanism that is different from the classical hand-over-hand mechanism of kinesin-1 (50, 53, 67). Further single-molecule work is needed to precisely determine how PAKRP2 achieves processive stepping and tight mechanochemical coupling despite having an extended neck linker domain. It is also not known how the long neck linker might impact the ability of PAKRP2 to generate sustained forces. From a biological perspective, there is still no direct evidence that PAKRP2 can associate with vesicle membranes and it is not clear whether intracellular transport by PAKRP2 generally results from a small population of motors or a large ensemble of motors attached to the cargo. Overall, these results add to the developing model of kinesin stepping, wherein each motor steps in a way that is optimized for its structure and role in the cell.

## Acknowledgments

This project was partially supported through grants from the National Science Foundation to W.Q. (1616462) and the National Institutes of Health to W.O.H. (R01GM076476). K.J.M was supported by a fellowship from the National Cancer Institute (F99CA223018).

## Author Contributions

W.Q conceived and supervised the study. A.G. and P.W. performed all experiments. K.J.M. built the microscope used for high-resolution tracking and assisted with experiments and analysis. A.G., W.O.H. and W.Q. wrote the manuscript with input from all authors.

---

**Figure S1:**
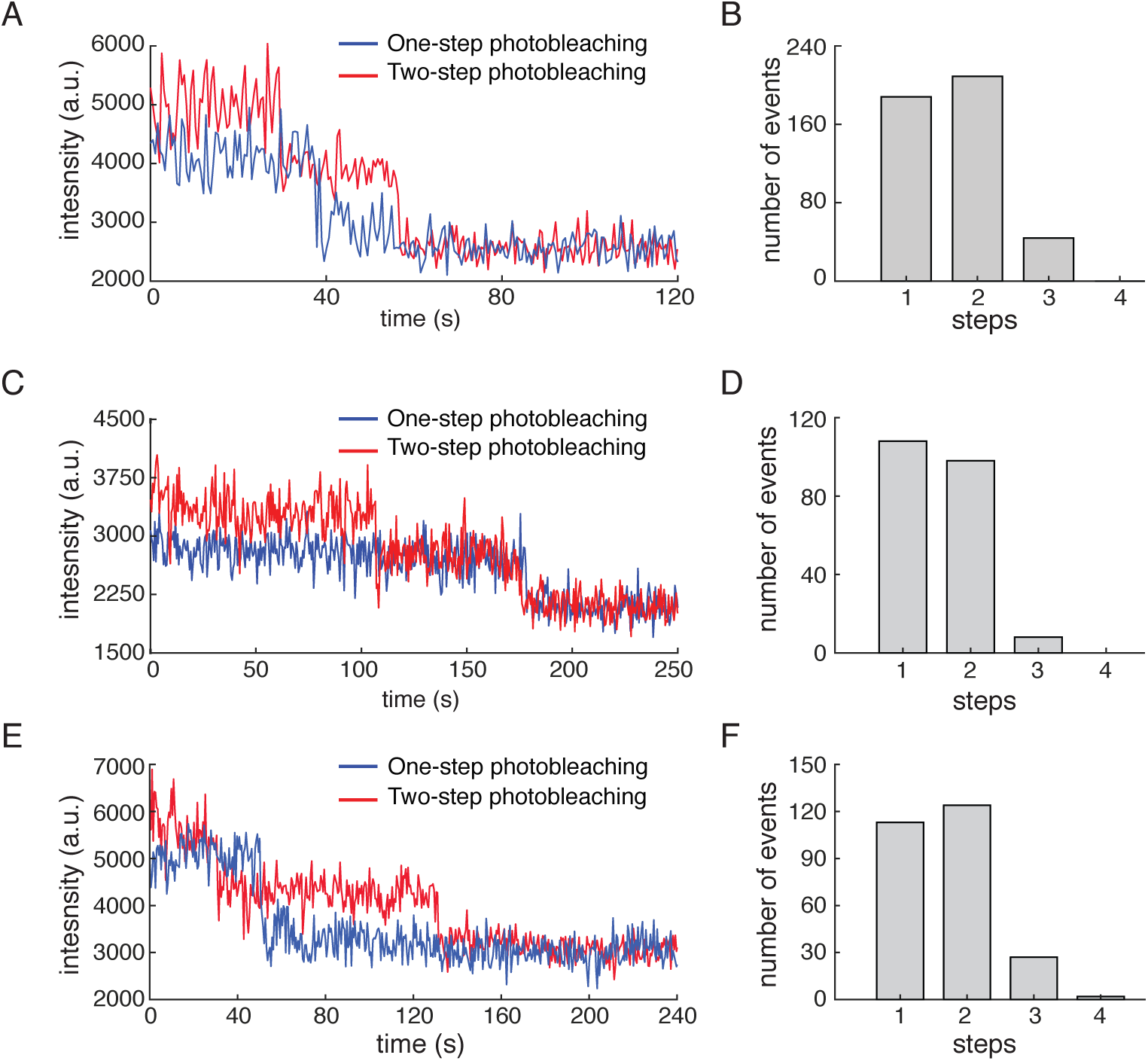
PAKRP2(FL), PAKRP2(560) and PAKRP2(LZ) form homodimers in solution. (A) Representative photobleaching traces of individual PAKRP2(FL) molecules. (B) Photobleaching histogram of PAKRP2(FL) (n = 441). (C) Representative photobleaching traces of individual PAKRP2(560) molecules. (D) Photobleaching histogram of PAKRP2(560) (n = 214). (E) Representative photobleaching traces of individual PAKRP2(LZ) molecules. (F) Photobleaching histogram of PAKRP2(LZ) (n = 266).

**Figure S2:**
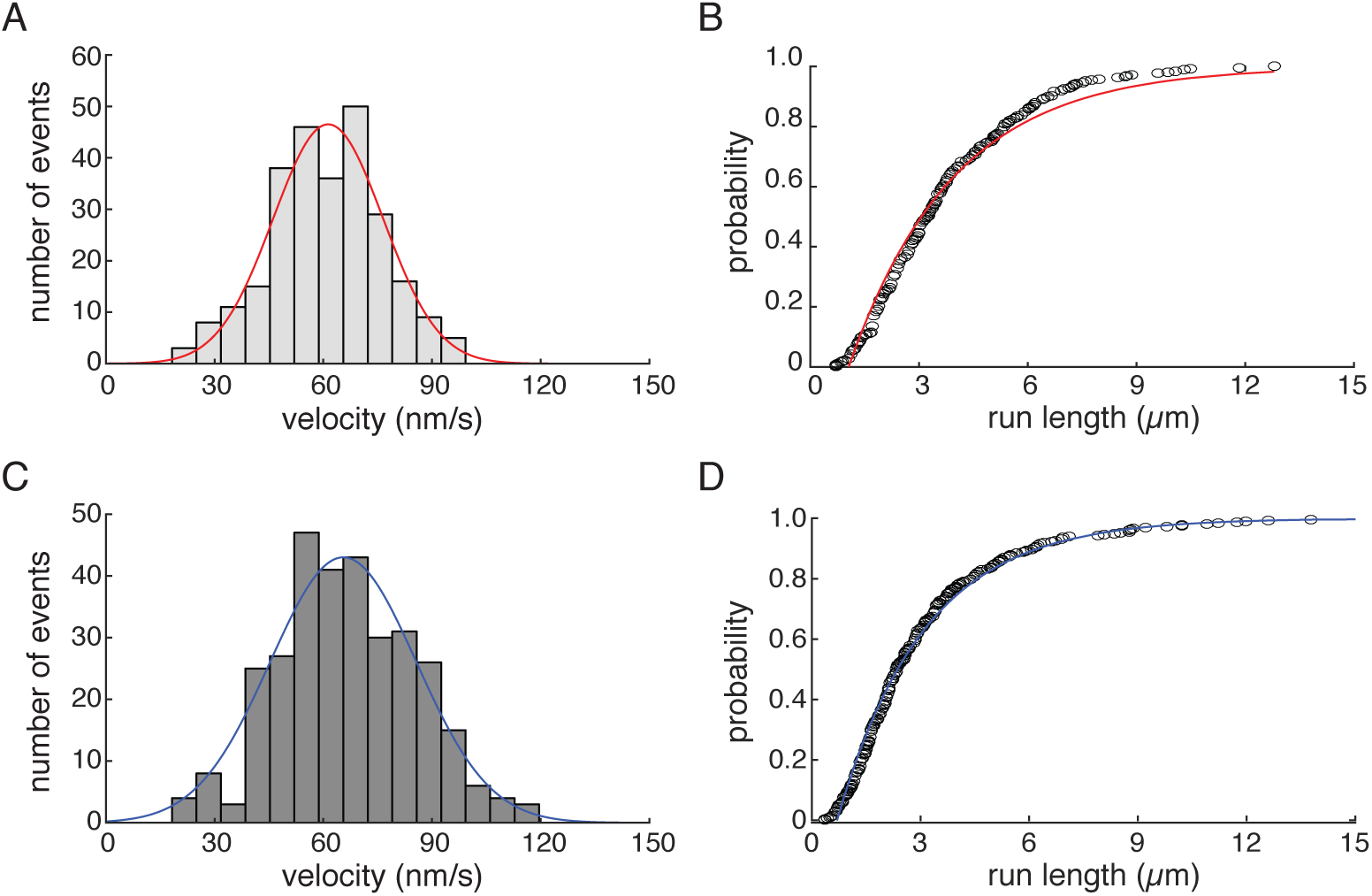
PAKRP2(560) and PAKRP2(LZ) velocity and run lengths. (A) Velocity histogram of PAKRP2(560). Red line corresponds to a Gaussian fit with a mean of 61 ± 15 nm s^−1^ (n = 266). (B) Run length histogram of PAKRP2(560). Red line corresponds to an exponential fit with a characteristic run length of 3.35 ± 0.29 *μ*m (n = 266). (C) Velocity histogram of PAKRP2(LZ). Blue line corresponds to a Gaussian fit with a mean of 66 ± 20 nm s^−1^ (n = 333). (D) Run length histogram of PAKRP2(LZ). Blue line corresponds to an exponential fit with a characteristic run length of 2.67 ± 0.25 *μ*m (n = 333).

**Figure S3:**
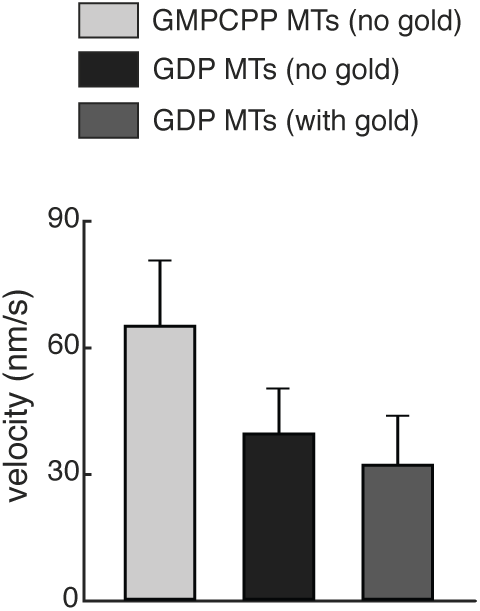
Gold nanoparticle does not affect motor activity. Bar graphs showing the relative velocities of PAKRP2(560) on GMPCPP or taxol-stabilized GDP microtubules without gold, and on taxol-stabilized GDP microtubules with gold. The velocities are 65 ± 16 nm s^−1^ (n = 271), 40 ± 12 nm s^−1^ (n = 47) and 32 ± 11 nm s^−1^ (n = 57), respectively.

**Figure S4:**
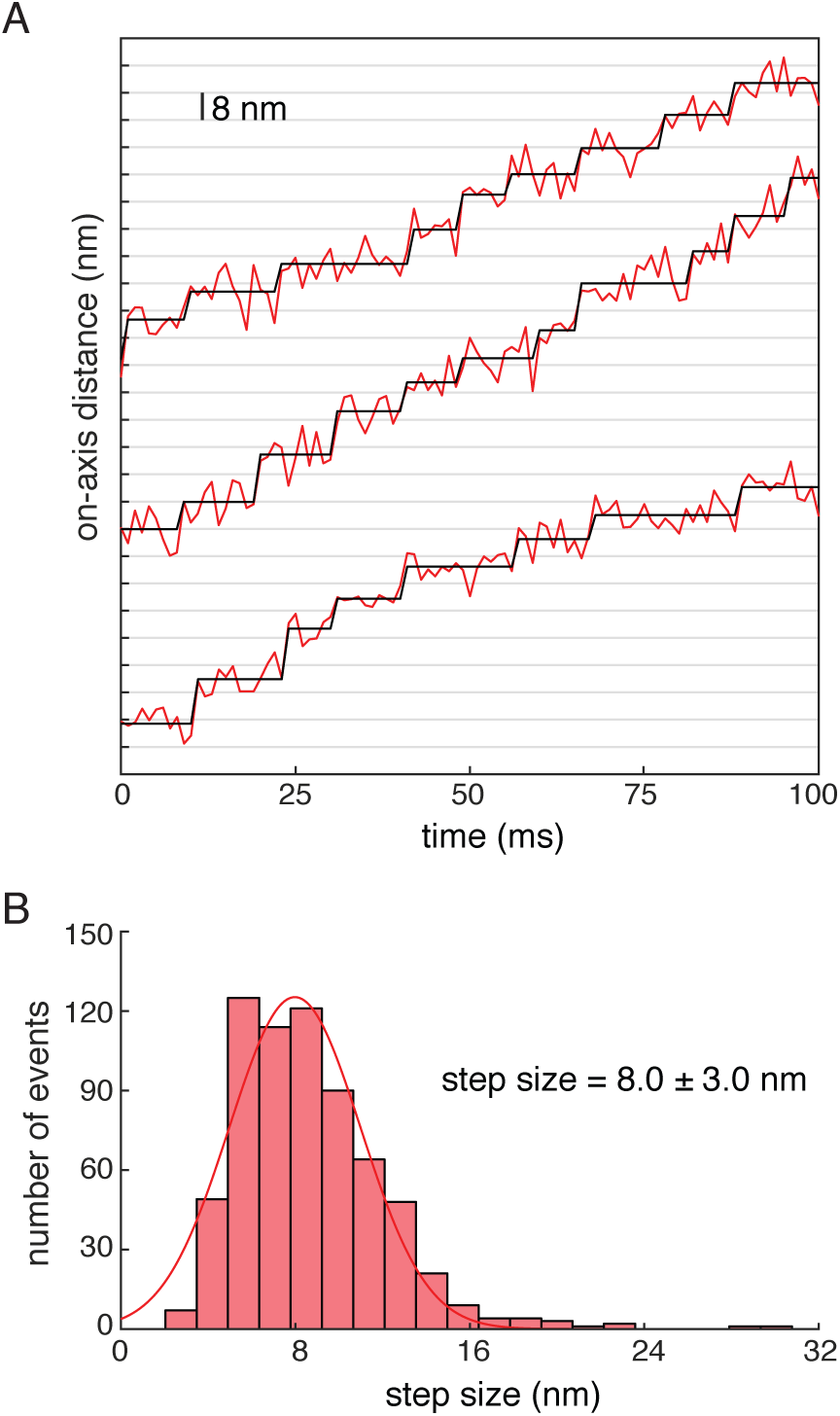
Step size determination for K560AviC. (A) A representative on-axis distance vs. time plot. Raw data is shown as red lines and steps detected by a step finding algorithm are shown in black. Data were acquired at 1 mM ATP every 1 ms. (B) Step size histogram for K560AviC with a single Gaussian fit (n = 664).

**Figure S5:**
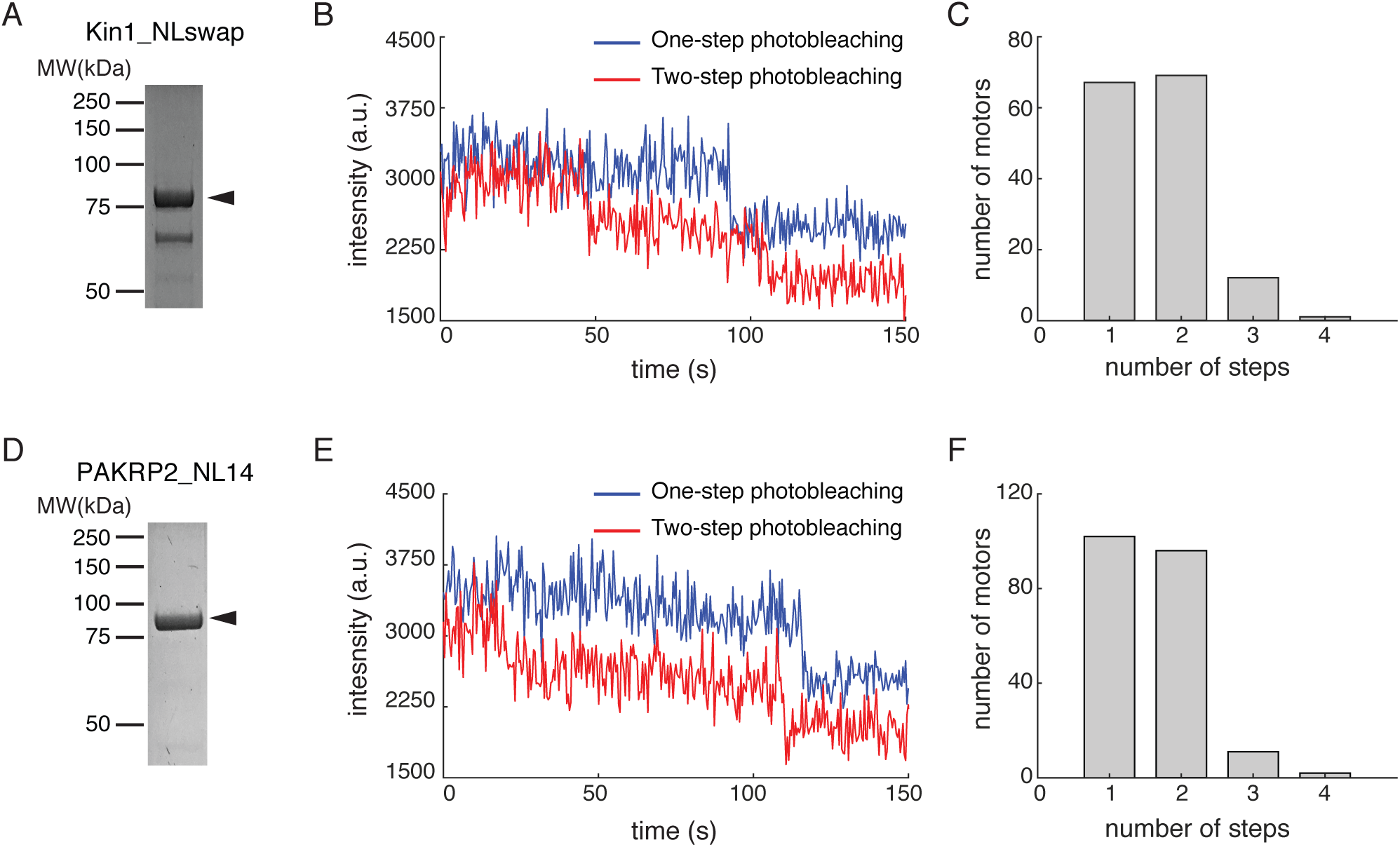
PAKRP2_NL14 and Kin1_NLswap both form individual homodimers in solution. (A) SDS-PAGE gel of Kin1_NLswap. Arrowhead indicates the expected band of Kin1_NLswap. MW = 75 kDa. (B) Representative photobleaching traces of individual Kin1_NLswap molecules. (C) Photobleaching histogram for Kin1_NLswap (n = 149). (D) SDS-PAGE gel of PAKRP2_NL14. Arrowhead indicates the expected band of PAKRP2_NL14. MW = 89 kDa. (E) Representative photobleaching traces of individual PAKRP2_NL14 molecules. (F) Photobleaching histogram for PAKRP2_NL14 (n = 289).

**Figure S6:**
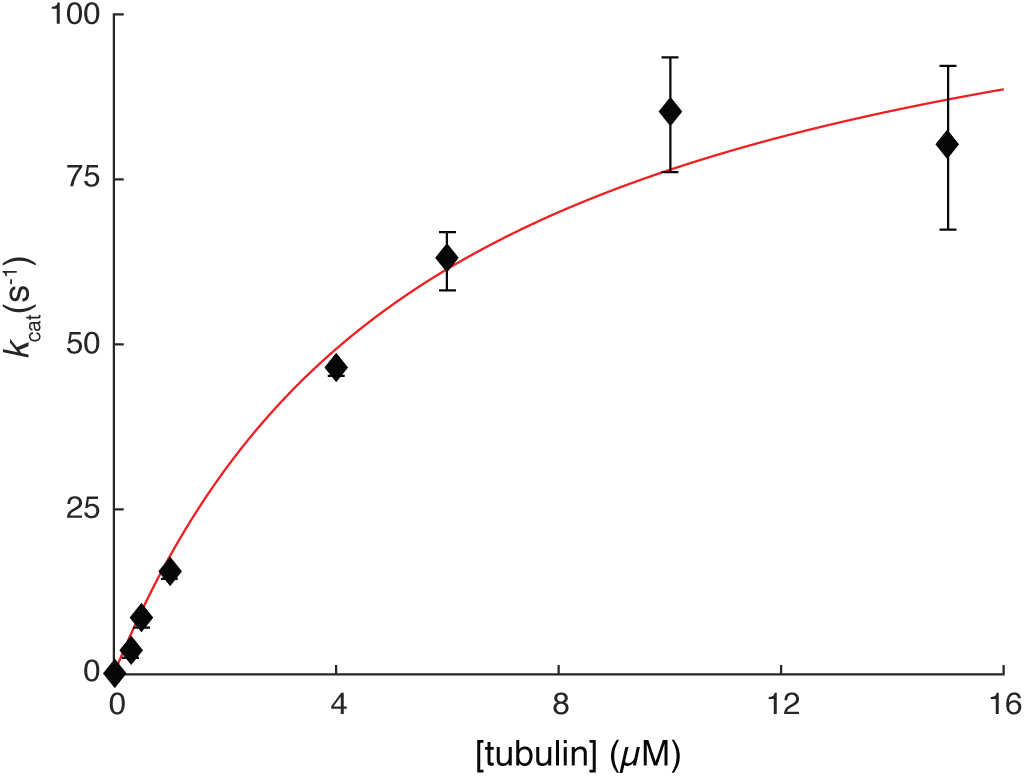
Kin_NLswap ATP hydrolysis and stepping rate are uncoupled. The microtubule-stimulated ATPase rate per dimer for Kin1_NLswap as a function of tubulin concentration. Data correspond to the mean ± SE (n = 3 determinations per point). The solid line represents a fit to the Michaelis-Menten equation, giving a maximum k_cat_ of 121 ± 7 s^−1^.

